## Supplemental Files for "Exploring the Pathogen Profiles of Ancient Feces"

**Table S1.** Primer and probe sequences for qPCR assays on the custom TAC

NOTE: Though the TAC contained RNA targets, only the DNA targets were used in this study

| **Pathogen** | **Gene** | **Primer or probe sequence (5' - 3')** | **Reference** |
| --- | --- | --- | --- |
| *Campylobacter jejuni*/*C. coli* | cadF | Fwd: CTGCTAAACCATAGAAATAAAATTTCTCAC |  |
|  |  | Rev: CTTTGAAGGTAATTTAGATATGGATAATCG | [1] |
|  |  | Probe: CATTTTGACGATTTTTGGCTTGA |  |
| *C. difficile* | tcdB | Fwd: GGTATTACCTAATGCTCCAAATAG |  |
|  |  | Rev: TTTGTGCCATCATTTTCTAAGC | [1] |
|  |  | Probe: CCTGGTGTCCATCCTGTTTC |  |
| EAEC (aaiC) | *aaiC* | Fwd: ATTGTCCTCAGGCATTTCAC |  |
|  |  | Rev: ACGACACCCCTGATAAACAA | [1] |
|  |  | Probe: TAGTGCATACTCATCATTTAAG |  |
| EAEC (aatA) | *aatA* | Fwd: CTGGCGAAAGACTGTATCAT |  |
|  |  | Rev: TTTTGCTTCATAAGCCGATAGA | [1] |
|  |  | Probe: TGGTTCTCATCTATTACAGACAGC |  |
| STEC (stx1) | *stx1* | Fwd: ACTTCTCGACTGCAAAGACGTATG |  |
|  |  | Rev: ACAAATTATCCCCTGWGCCACTATC | [1] |
|  |  | Probe: 56FAM/CTCTGCAATAGGTACTCCA/3MGB-NFQ/ |  |
| STEC (stx2) | *stx2* | F, CCACATCGGTGTCTGTTATTAACC |  |
|  |  | R, GGTCAAAACGCGCCTGATAG | [1] |
|  |  | P, 5VIC/TTGCTGTGGATATACGAGG/3MGB-NFQ/ |  |
| EPEC (eae) | *eae* | F, CATTGATCAGGATTTTTCTGGTGATA |  |
|  |  | R, CTCATGCGGAAATAGCCGTTA | [1] |
|  |  | P, 56FAM/ATACTGGCGAGACTATTTCAA/3MGB-NFQ/ |  |
| EPEC (bfpA) | *bfpA* | F, TGGTGCTTGCGCTTGCT |  |
|  |  | R, CGTTGCGCTCATTACTTCTG | [1] |
|  |  | P, 5VIC/CAGTCTGCGTCTGATTCCAA/3MGB-NFQ/ |  |
| ETEC LT | *LT* | F, TTCCCACCGGATCACCAA |  |
|  |  | R, CAACCTTGTGGTGCATGATGA | [1] |
|  |  | P, CTTGGAGAGAAGAACCCT |  |
| ETEC ST | *ST* | Fh, GCTAAACCAGYAGRGTCTTCAAAA |  |
|  |  | Fp, TGAATCACTTGACTCTTCAAAA |  |
|  |  | Rh, CCCGGTACARGCAGGATTACAACA | [1] |
|  |  | Rp, GGCAGGATTACAACAAAGTT |  |
|  |  | Ph, 6VIC/TGGTCCTGAAAGCATGAA/3MGB-NFQ/ |  |
|  |  | Pp, 6VIC/TGAACAACACATTTTACTGCT/3MGB-NFQ/ |  |
| EIEC/*Shigella* | *ipaH* | F, CCTTTTCCGCGTTCCTTGA |  |
|  |  | R, CGGAATCCGGAGGTATTGC | [1] |
|  |  | P, 56FAM/CGCCTTTCCGATACCGTCTCTGCA/3MGB-NFQ/ |  |
| *Salmonella* | *ttr* | Fwd: CTCACCAGGAGATTACAACATGG |  |
|  |  | Rev: AGCTCAGACCAAAAGTGACCATC | [1] |
|  | MGB probe | Probe: CACCGACGGCGAGACCGACTTT |  |
| *E. coli* O157 | *rfbE* | Fwd: TTTCACACTTATTGGATGGTCTCAA |  |
|  |  | Rev: CGATGAGTTTATCTGCAAGGTGAT | [1] |
|  |  | Probe: CTCTCTTTCCTCTGCGGTCCT |  |
| *Cryptosporidium* | *18S* | Fwd: GGGTTGTATTTATTAGATAAAGAACCA |  |
|  |  | Rev: AGGCCAATACCCTACCGTCT | [1] |
|  |  | Probe: TGACATATCATTCAAGTTTCTGAC |  |
| *Giardia* spp. | *18S* | Fwd: GACGGCTCAGGACAACGGTT |  |
|  |  | Rev: TTGCCAGCGGTGTCCG | [1] |
|  |  | Probe: CCCGCGGCGGTCCCTGCTAG |  |
| *E. histolytica* | *18S* | Fwd: ATTGTCGTGGCATCCTAACTCA |  |
|  |  | Rev: GCGGACGGCTCATTATAACA | [1] |
|  |  | Probe: TCATTGAATGAATTGGCCATTT |  |
| *Entamoeba* spp. | *18S rRNA* | Fwd: AAACGATGTCAACCAAGGATTG |  |
|  |  | Rev: TCCCCCTGAAGTCCATAAACTC | [1] |
|  |  | Probe: CCTTGTTCAGAACTTAAAGAGAAA |  |
| *Ascaris* | *ITS1* | Fwd: GCCACATAGTAAATTGCACACAAAT |  |
|  |  | Rev: GCCTTTCTAACAAGCCCAACAT | [1] |
|  |  | Probe: TTGGCGGACAATTGCATGCGAT |  |
| *Trichuris* | *18S rRNA* | Fwd: TTGAAACGACTTGCTCATCAACTT |  |
|  |  | Rev: CTGATTCTCCGTTAACCGTTGTC | [1] |
|  |  | Probe: CGATGGTACGCTACGTGCTTACCATGG |  |
| *Necator americanus* | ITS-2 | Fwd: CTGTTTGTCGAACGGTACTTGC |  |
|  |  | Rev: ATAACAGCGTGCACATGTTGC | [1] |
|  |  | Probe: CTGTACTACGCATTGTATAC |  |
| *Strongyloides stercoralis* | dispered repetitive sequence | Fwd: TCCAGAAAAGTCTTCACTCTCCAG |  |
|  |  | Rev: TGCGTTAGAATTTAGATATTATTGTTGCT | [1] |
|  |  | Probe: TCAGCTCCAGTTGAACAACAGCCTCCAA |  |
| *Blastocystis* spp. | 18s rRNA | Fwd: TGGTCCGRTGAACACTTTGGAT |  |
|  |  | Rev: CCTACGGAAACCTTGTTACGACTTCA | [1] |
|  |  | Probe: CTTCCTCTAAATGRTAAGATT |  |
| *Ancylostoma duodenales* | ITS-2 | Fwd: GAATGACAGCAAACTCGTTGTTG |  |
|  |  | Rev: ATACTAGCCACTGCCGAAACGT | [1] |
|  |  | Probe: ATCGTTTACCGACTTTAG |  |
| *Enterobius vermicularis* | 5S rRNA | Fwd: CAAACAACTGCATCACCAATAAC |  |
|  |  | Rev: AGTGTAGAGCAATAAGCAGTAAAG | [2] |
|  |  | Probe: TACCAACAACACTTGCACGTCTCTTCA |  |
| *H. nana* | ITS1 | Fwd: CATTGTGTACCAAATTGATGATGAGTA |  |
|  |  | Rev: CAACTGACAGCATGTTTCGATATG | [1] |
|  |  | Probe: CGTGTGCGCCTCTGGCTTACCG |  |
| enteric 16s |  | Fwd: TGCAAGTCGAACGAAGCACTTTA |  |
|  |  | Rev: GCAGGTTACCCACGCGTTAC | [1] |
|  |  | Probe: CGCCACTCAGTCACAAA |  |
| PhHV | gB | Fwd: GGGCGAATCACAGATTGAATC |  |
|  |  | Rev: GCGGTTCCAAACGTACCAA | [1] |
|  |  | Probe: TATGTGTCCGCCACCATCT |  |
| *Yersinia enterocolitica* | *lytA* | Fwd: TGATTCACCAGCAGCAATAC |  |
|  |  | Rev: GGCATCATGAAAGGCGG | [1] |
|  |  | Probe: TGTCGGTTTCTCCTTCCAGG |  |
| *Heliobacter pylori* | *ureC* | Fwd: GACACCAGAAAAAGCGGCTA |  |
|  |  | Rev: AGCGCATGTCTTCGGTTAAA | [1] |
|  |  | Probe: TCACTAAAGCGTTTTCTACC |  |
| *Plesiomonas shigelloides* | *gyrB* | Fwd: CCGCCGTGAAGGCAAAG |  |
|  |  | Rev: GCTACCGGCTCACCCAGAT | [1] |
|  |  | Probe: CACACCCAAGAATAC |  |
| *Cyclospora cayetanensi* | 18s rRNA | Fwd: AAAAGCTCGTAGTTGGATTTCTG |  |
|  |  | Rev: AACACCAACGCACGCAGC | [1] |
|  |  | Probe: AAGGCCGGATGACCACGA |  |
| *Cystoisospora belli* | 18s rRNA | Fwd: ATATTCCCTGCAGCATGTCTGTTT |  |
|  |  | Rev: CCACACGCGTATTCCAGAGA | [1] |
|  |  | Probe: CAAGTTCTGCTCACGCGCTTCTGG |  |
| *Blastocystis* spp. | 18s rRNA | Fwd: TGGTCCGRTGAACACTTTGGAT |  |
|  |  | Rev: CCTACGGAAACCTTGTTACGACTTCA | [1] |
|  |  | Probe: CTTCCTCTAAATGRTAAGATT |  |
| *Enterocytozoon bieneusi* | SSU rRNA | Fwd: TGTGTAGGCGTGAGAGTGTATCTG |  |
|  |  | Rev: CATCCAACCATCACGTACCAATC | [1] |
|  |  | Probe: CACTGCACCCACATCCCTCACCCTT |  |
| *Encephalitozoon intestinalis* | ITS | Fwd: CACCAGGTTGATTCTGCCTGAC |  |
|  |  | Rev: CTAGTTAGGCCATTACCCTAACTACCA | [1] |
|  |  | Probe: CTATCACTGAGCCGTCC |  |
| *Balantidium coli* | ITS-1 | Fwd: TGCAATGTGAATTGCAGAACC |  |
|  |  | Rev: TGGTTACGCACACTGAAACAA | [1] |
|  |  | Probe: CTGGTTTAGCCAGTGCCAGTTGC |  |
| *Acanthamoeba* spp. | 18S rRNA | Fwd: CCCAGATCGTTTACCGTGAA |  |
|  |  | Rev: TAAATATTAATGCCCCCAACTATC | [4] |
|  |  | Probe: CTGCCACCGAATACATTAGCATGG |  |

**Table S2.** TaqMan Array Card (TAC) performance and standard curve parameters**.**

| **Target** | **Target Gene** | **Slope** | **Y-intercept** | **R^2^** | **Efficiency** | **95% LOD† (copies/µL template)** |
| --- | --- | --- | --- | --- | --- | --- |
| enteric 16S | 16S | -3.309 | 38.881 | 0.998 | 101% | 1 |
| *Acanthamoeba* spp. | 18S rRNA | -3.3877 | 37.82 | 1.000 | 97% | 38 |
| *Ancylostoma duodenale* | ITS-2 | -3.3832 | 39.101 | 1.000 | 98% | 10 |
| *Ascaris lumbricoides* | ITS-1 | -3.4482 | 38.594 | 1.000 | 95% | 10 |
| *Balantidium coli* | ITS-1 | -3.3935 | 37.92 | 1.000 | 97% | 4 |
| *Blastocystis* spp. | 18S rRNA | -3.3196 | 40.64 | 0.997 | 100% | 4 |
| *Cystoisospora belli* | 18S rRNA | -3.3479 | 37.801 | 0.999 | 99% | 10 |
| *Cyclospora cayetanensi* | 18S rRNA | -3.3408 | 37.151 | 0.998 | 99% | 4 |
| *Campylobacter jejuni/coli* | *cadF* | -3.34178 | 38.27 | 0.999 | 99% | 35 |
| *Clostridium difficile* | *tcdB* | -3.4282 | 37.542 | 0.999 | 96% | 10 |
| *Cryptosporidium* spp. | 18S rRNA | -3.4033 | 37.983 | 0.999 | 97% | 1 |
| DNA control (phocine herpes virus) | *gB* | -3.315 | 37.009 | 0.998 | 100% | 10 |
| *Enterocytozoon bieneusi* | ITS | -3.2802 | 37.209 | 0.999 | 102% | 8 |
| *E. coli* O157:H7 | *rfbE* | -3.4568 | 37.976 | 1.000 | 95% | 4 |
| *Encephalitozoon intestinalis* | SSU rRNA | -3.3819 | 38.462 | 0.999 | 98% | 4 |
| *Enterobius vermicularis* | 5S | -3.4592 | 38.572 | 0.999 | 95% | 120 |
| EAEC (aaiC) | *aaiC* | -3.4241 | 38.15 | 0.999 | 96% | 10 |
| EAEC (aatA) | *aatA* | -3.4252 | 37.694 | 0.998 | 96% | 38 |
| *Entamoeba hystolytica* | 18S rRNA | -3.2775 | 37.994 | 0.996 | 102% | 10 |
| *Entamoeba* spp. | 18S rRNA | -3.2317 | 37.259 | 0.974 | 104% | 35 |
| EPEC (typical) | *bfpA* | -3.3772 | 37.465 | 0.999 | 98% | 10 |
| EPEC (atypical) | *eae* | -3.372 | 37.592 | 0.999 | 98% | 4 |
| ETEC (LT) | *LT* | -3.4638 | 47.637 | 0.990 | 94% | 485 |
| ETEC (STh) | *STh* | -3.3785 | 38.763 | 0.999 | 98% | 10 |
| ETEC (STp) | *STp* | -3.3548 | 37.266 | 0.999 | 99% | 4 |
| *Giardia* spp. | 18S rRNA | -3.4182 | 37.863 | 1.000 | 96% | 10 |
| *Hymenolepis nana* |  | -3.3804 | 38.248 | 1.000 | 98% | 4 |
| *Helicobacter pylori* | *ureC* | -3.4078 | 37.726 | 0.998 | 97% | 10 |
| *Shigella*/EIEC | *ipaH* | -3.3522 | 37.506 | 0.999 | 99% | 38 |
| *Necator americanus* | ITS-2 | -3.3686 | 39.806 | 1.000 | 98% | 7 |
| *Plesiomonas shigelloides* | *gyrB* | -3.4184 | 38.202 | 1.000 | 96% | 38 |
| *Salmonella* spp. | *invA* | -3.4191 | 38.427 | 1.000 | 96% | 4 |
| *Strongyloides stercolaris* | Dispersed repetitive sequence | -3.3316 | 37.527 | 0.999 | 100% | 4 |
| STEC (stx1) | *stx1* | -3.4083 | 39.883 | 1.000 | 97% | 120 |
| STEC (stx2) | *stx2* | -3.3697 | 38.328 | 0.967 | 98% | 160 |
| *Trichuris trichiura* | 18S rRNA | -3.3502 | 38.395 | 1.000 | 99% | 4 |
| *Yersinia enterocolitica* | *lytA* | -3.483 | 38.279 | 0.998 | 94% | 4 |

†95% LOD in gene copies per reaction calculated using methods from Stokdyk *et al*. 2016 [6]

**Figure S1. Amplification and multicomponent plots**

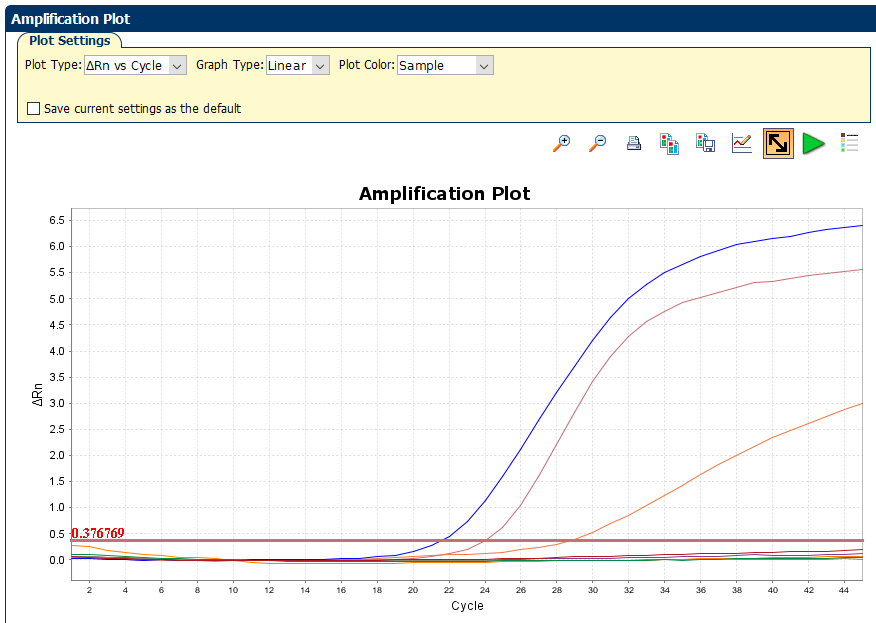

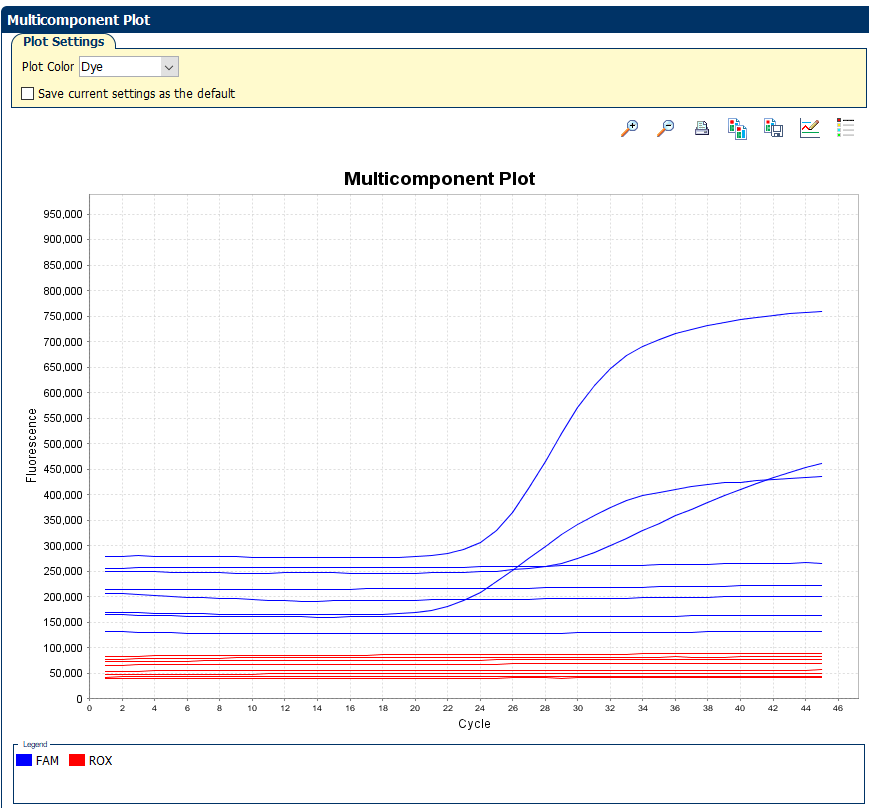

Table S3. MIQE Checklist

| **ITEM TO CHECK** | **IMPORTANCE** | **CHECKLIST** |
| --- | --- | --- |
| **EXPERIMENTAL DESIGN** |  |  |
| Definition of experimental and control groups | **E** | Cross-sectional study with no intervention or control group |
| Number within each group | **E** | 10 paleofeces from Mexico and 12 from Oregon |
| Assay carried out by core lab or investigator's lab? | D | Investigator's lab |
| **SAMPLE** |  |  |
| Description | **E** | 200 mg (see method’s section) |
| Volume/mass of sample processed | D | 200 mg |
| Microdissection or macrodissection | **E** | Not applicable |
| Processing procedure | **E** | See method’s section. |
| If frozen - how and how quickly? | **E** | Not frozen |
| If fixed - with what, how quickly? | **E** | Not fixed |
| Sample storage conditions and duration (especially for FFPE samples) | **E** | Shipped at ambient conditions. Stored at room temperature away a dark, cool, dry place |
| **NUCLEIC ACID EXTRACTION** |  |  |
| Procedure and/or instrumentation | **E** | See methods section |
| Name of kit and details of any modifications | **E** | Adapted from Hagan et al. 2020 [7] |
| Source of additional reagents used | D | Qiagen PowerBead Tubes with garnet beads |
| Details of DNase or RNAse treatment | **E** | Not applicable |
| Contamination assessment (DNA or RNA) | **E** | One extraction negative control was included during each day of extractions |
| Nucleic acid quantification | **E** | Qubit 1X HS dsDNA Kit |
| Instrument and method | **E** | Qubit 4 Fluorometer |
| RNA integrity method/instrument | **E** | Not measured |
| Inhibition testing (Cq dilutions, spike or other) | **E** | Monitored amplification of spiked controls |
| **qPCR TARGET INFORMATION** |  |  |
| If multiplex, efficiency and LOD of each assay. | **E** | Table S2 |
| *In silico* specificity screen (BLAST, etc) | **E** | We BLASTed all assays to confirm specificity before ordering the custom TAC. |
| **qPCR OLIGONUCLEOTIDES** |  |  |
| Primer sequences | **E** | Table S1 |
| Probe sequences | D** | Table S1 |
| Location and identity of any modifications | **E** | No modifications |
| Manufacturer of oligonucleotides | D | ThermoFisher Scientific |
| **qPCR PROTOCOL** |  |  |
| Complete reaction conditions | **E** | 45°C for 20 min and 95°C for 10 min, followed by 45 cycles of 95°C for 15 s and 60°C for 1 min |
| Reaction volume and amount of cDNA/DNA | **E** | 6.66 µL of template, 31.33 µL molecular grade water, 2 µL inhibition control, with 60 µL of AgPath-ID™ One-Step RT-PCR Reagents |
| Primer, (probe), Mg++ and dNTP concentrations | **E** | All assays contained the same concentrations of primers (900 nanomolar) and probe (250 nanomolar). The Mg2+ and dNTP concentrations are not listed in the in the User Guide. |
| Polymerase identity and concentration | **E** | AmpliTaq Gold™ polymerase |
| Buffer/kit identity and manufacturer | **E** | AgPath-ID™ One-Step RT-PCR Reagents |
| Additives (SYBR Green I, DMSO, etc.) | **E** | No additives |
| Manufacturer of plates/tubes and catalog number | D | ThermoFisher Scientific |
| Complete thermocycling parameters | **E** | 45°C for 20 min and 95°C for 10 min, followed by 45 cycles of 95°C for 15 s and 60°C for 1 min |
| Reaction setup (manual/robotic) | D | Manual set-up in a disinfected dead air box (10% bleach with fifteen minutes of contact time, UV for fifteen minutes, and a final cleaning step with 70% ethanol) |
| Manufacturer of qPCR instrument | **E** | ThermoFisher Scientfic |
| **qPCR VALIDATION** |  |  |
| Evidence of optimisation (from gradients) | D | See Liu *et al*. 2013 [8] and Liu *et al*. 2016 [9] |
| Specificity (gel, sequence, melt, or digest) | **E** | See Liu *et al*. 2013 [8] and Liu *et al*. 2016 [9] |
| Standard curves with slope and y-intercept | **E** | Table S2 |
| PCR efficiency calculated from slope | **E** | Table S2 |
| r2 of standard curve | **E** | Table S2 |
| Evidence for limit of detection | **E** | Table S2 |
| **DATA ANALYSIS** |  |  |
| qPCR analysis program (source, version) | **E** | QuantStudio Real-Time PCR Software V1.2 CDC |
| Cq method determination | **E** | Manual thresholding |
| Results of NTCs | **E** | Reported in the results section |
| Justification of number and choice of reference genes | **E** | N/A |
| Description of normalisation method | **E** | Normalized to mass of paleofeces |
| Software (source, version) | E | R Studio V2.2.2 |

Table S4. hCYTB484 human mtDNA primers and probe sequences.

| Oligonucleotide | Sequence (5’ to 3’) | Reference |
| --- | --- | --- |
| Fwd primer | CAATGAATCTGAGGAGGCTAC | Zhu, K.; Suttner, B.; Pickering, A.; Konstantinidis, K. T.; Brown, J. A Novel Droplet Digital PCR  Human MtDNA Assay for Fecal Source Tracking. Water Res. 2020, 183, 116085.  https://doi.org/10.1016/J.WATRES.2020.116085. |
| Rev primer | CGTGCAAGAATAGGAGGTG |  |
| Probe | ACCCTCACACGATTCTTTACCTTTCACT |  |
